# Lower childhood subjective social status is associated with greater neural responses to ambient auditory deviance

**DOI:** 10.1101/2022.12.22.521644

**Authors:** Yu Hao, Lingyan Hu

## Abstract

Humans’ early life experience varies by socioeconomic status, raising the question of how this difference is reflected in the adult brain. An important aspect of brain function is the ability to detect salient ambient changes while focusing on a task. Here we ask whether perceived childhood social standing is reflected by the way young adults’ brain signals correlate with detecting changes in irrelevant information. In two studies (total N = 58), we examine electrical brain responses in the frontocentral region to a series of auditory tones, consisting of “standard” stimuli (80%) and “deviant” stimuli (20%) interspersed randomly, while participants were engaged in various visual tasks. Both studies showed stronger automatic change detection indexed by mismatch negativity (MMN) in lower-SES individuals, regardless of the sound’s features, emotional content, or study type. Moreover, we observed a larger MMN in lower-SES participants, even though they did not show differences in brain and behavior responses to the attended task and did not involuntarily orient more attention to deviance, as indexed by the P3a. The study indicates that individuals with lower subjective social status may have an increased ability to automatically detect changes in their environment, which may suggest their adaptation to their childhood environments.

## Introduction

Socioeconomic status (SES) describes a person’s resources, such as income, education, neighborhood quality, as well as their perceived social standing (Farah, 2017). Studies have demonstrated that disparities in SES, including parental education and income or retrospectively reported childhood poverty, can have an impact on cognitive and socioemotional development, as well as brain structure and function, in both children and adults (Farah, 2017; Letourneau et al., 2013; Noble & Giebler, 2020).

One area of study regarding the impact of SES on cognitive outcomes is task-irrelevant information processing. Previous research has shown that lower-SES children tend to be more responsive to irrelevant information that they were supposed to disregard compared to their higher-SES peers. For instance, they tended to perform worse when distractions were present, such as slower reaction time in incongruent trials in the Flanker task (Mezzacappa, 2004), recognized more distracting, irrelevant images in reading materials (Norman and Breznitz, 1992), and chose a lower signal-to-noise ratio when listening to a speaker in a noisy background (Evans et al., 1995). These outcomes were traditionally seen as a deficit in lower-SES children’s concentration skills. However, a recent adaptation-based perspective suggests that the greater attention to distractions in lower-SES children could be an adaptive response to harsh living environments (e.g., Ellis et al., 2017; Frankenhuis et al., 2020). More likely to live in unpredictable and stressful environments, lower-SES children may benefit from staying vigilant against unpredictable physical or social threats, which could lead to this increased sensitivity to task-irrelevant information.

To explore SES-related differences in neural responses to task-irrelevant information, some studies have used event-related potential (ERP), which measures very small voltages generated in the brain in response to specific events or stimuli. These ERP studies have used selective attention paradigms, where participants are instructed to either attend to sounds in only one ear while sound inputs were played binaurally (e.g., Stevens et al., 2009), or respond to specific types of tones while ignoring others in a sound sequence (e.g., D’Angiulli et al., 2008). Interestingly, lower-SES children appeared to show attenuated selective attention, as reflected by reduced differences in ERP amplitudes evoked by attended vs. unattended stimuli, but without an observable performance gap (D’Angiulli et al., 2008; D’Angiulli, Weinberg, et al., 2012; Li et al., 2022; Stevens et al., 2009, 2015). In particular, Steven et al. (2009) found that lower-SES children had similar ERP amplitudes as their higher-SES peers to attended probes but stronger ERP responses to the unattended ones. D’Angiulli et al. (2008; 2012) found that lower-SES children showed higher theta power to the unattended tones than the attended one. These findings suggest that lower-SES children may tend not to suppress task-irrelevant information, which some researchers interpret as an adaptation to living in more unpredictable environments, rather than a deficit in attention (D’Angiulli, Lipina, et al., 2012).

The aim of the present study is to contribute to the existing literature on how SES affects cognitive processing of task-irrelevant information. Specifically, we focus on a particular component of ERP, mismatch negativity (MMN), in a group of young adults. MMN is generated when there is a deviation in a sequence of auditory stimuli, and it peaks at around 100-250ms after the change onset (Näätänen et al., 2007; Näätänen & Kreegipuu, 2011). Unlike other ERP studies summarized above, MMN is automatic and occurs without effortful control of attention (Näätänen et al., 2007; Näätänen & Kreegipuu, 2011; Sussman et al., 2014). It is also reliable and can be used to indicate individual differences in cognitive functioning (e.g., Light et al. 2007; Ding et al. 2022). Our study uses a passive-listening paradigm, where participants were instructed to focus on visual information while we recorded their MMN in response to changes in auditory stimuli.

In addition, we aim to examine the impact of subjective social status during childhood, as retrospectively reported by the participants, on brain function in adulthood. While numerous studies have explored how objective measures of SES, such as income, job prestige, or education, affect brain structure and function (Noble & Giebler, 2020), these measures may not fully reflect how individuals perceived their social status. Our study uses the MacArthur Scale of Subjective Social Status (Adler et al., 2000), which may better tap into people’s feelings about their social status and thus have a closer connection to their judgments, emotional reactions and behaviors (Greitemeyer & Sagioglou, 2019). This measure of perceived social standing has been found to be an important predictor of later health outcomes and psychological functioning in a large body of interdisciplinary research, even when controlling for objective measures of SES (e.g., Adler et al. 2000a; Singh-Manoux et al. 2005; Hoebel et al. 2017; Anwyl-Irvine et al. 2021). Several neural studies have also shown unique associations when subjective social status and brain structure and function, independent of objective SES (Assari et al., 2020; Gianaros et al., 2008; Sheridan et al., 2013).

To build on previous research suggesting that lower-SES children are more sensitive to task-irrelevant information, we hypothesized that young adults with lower childhood subjective social status would exhibit greater brain responses to auditory change detection, as reflected by larger MMN amplitudes. We also examined the P3a ERP component, which reflects involuntary attention orientation to unattended sound changes (Friedman et al., 2001), to investigate whether lower childhood subjective social status was associated with a higher level of attention distraction. Additionally, we manipulated the attended visual information and salience of the unattended sound to explore whether lower-SES individuals showed a larger MMN in different contexts, given their tendency to exhibit a greater brain response to negative stimuli (Hao & Farah 2020).

## Methods

### Participants

Participants were undergraduate and graduate students from Cornell University. Exclusion criteria were medication use that could affect the nervous system and any neurological or psychiatric illness history. For Study 1, data was collected from 20 participants, but four did not report their socio-economic status (SES), resulting in a valid sample size of 16 (13 women, age = 19.94, S.D. = 2.32).

Based on the effect size of Study 1, we conducted a simulation-based power analysis (Kumle et al., 2021) for Study 2. The result from 1,000 simulations suggested that at the significance level of 0.05, a sample size of 40 is needed to reach a power of 95.9% (95% CI = 94.48 to 97.04). Therefore, data for Study 2 was collected from 50 participants, but eight were excluded: one participant fell asleep during recording, four did not report their SES, and three did not show clear MMN waveform. The final sample size was 42 (18 women, age = 21.93, S.D. = 3.79).

Table 1 presents the demographic information for both studies. The study was approved by the Cornell University Institutional Review Board (Protocol ID#: 1610006664), and participants provided informed consent. They were compensated with either course credits or $20.

**Table 1.** Participants’ demographic information.

| Sample | Study 1 (N = 16) | Study 2 (N=42) |
| --- | --- | --- |
| Gender<br>(women) | 81.3% | 66.7% |
| Age | M = 19.94<br>SD = 2.32 | M = 21.93<br>SD = 3.79 |
| Race | Asian: 8 or 50%<br>White: 6 or 37.5%<br>African American: 2 or 12.5% | Asian: 33 or 78.6%<br>White: 9 or 21.4%<br>African American: 0 or 0% |

### SES measurement

The participants’ SES was measured using the MacArthur Scale of Subjective Social Status, which asks participants to rank their family’s social standing during their childhood relative to their immediate community using a nine-rung “social ladder” (Figure 1). The question was asked: “Think of this ladder as representing where people stand in their communities. People define community in different ways; please define it in whatever way is most meaningful to you. At the top of the ladder are people who have the highest standing in their community. At the bottom are the people who have the lowest standing in their community. Where would you place your family when you were 8-10 years old on this ladder? Please place a large “X” on the rung where you think you stand at this time in your life relative to other people in your community.” This comparison was considered more relevant to their perceived social standing than comparing to the entire country where they grew up, as it reflects their proximate environment and social interactions (Cundiff et al. 2013). Higher scores indicate higher childhood family social standing, ranging from 1 to 9.

**Figure 1.**
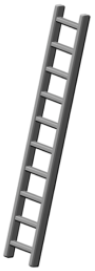
MacArthur ladder of Subjective Social Status.

Since many participants were international students, parental income or educational attainment measures were not comparable, and most participants considered themselves affluent in material resources (measured by perceived financial difficulties and neighborhood resources, Griskevicius et al., 2011, see Figure 2 (a-b)). However, there was considerable variability in their perceived social standing (Figure 2 (c)), with mean scores of 6.63 (S.D. = 1.39) in Study 1 and (S.D. = 1.44) in Study 2. The SES was not associated with age (*p* = 0.737), gender (*p* = 0.680), or race (*p* = 0.200) in the combined sample of 58 participants.

**Figure 2.**
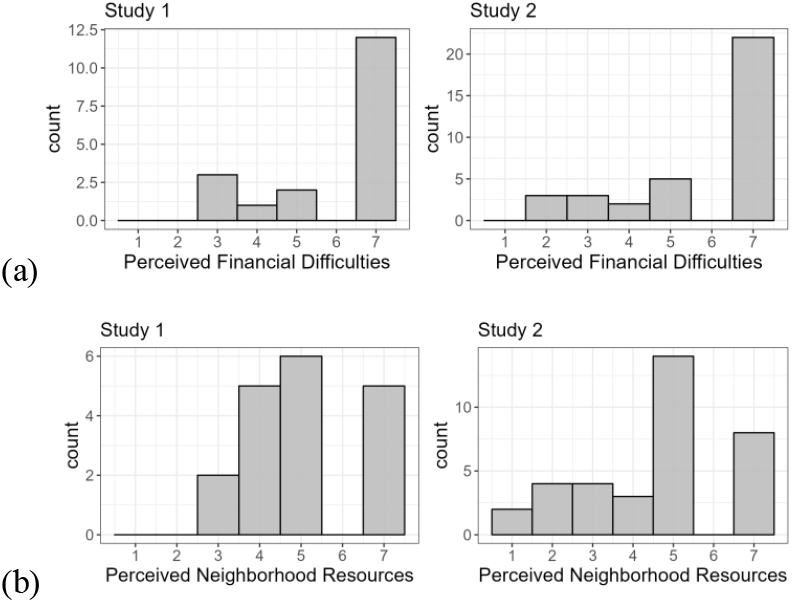
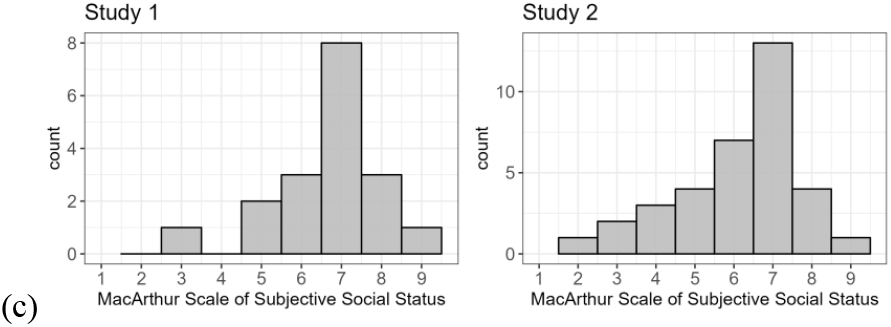
Histogram of the Perceived Financial Difficulties scale and the Perceived Neighborhood Resources (a-b) and histogram of the MacArthur Scale of Subjective Social Status.

Perceived Stress Scale (PSS; Cohen, Kamarck, & Mermelstein, 1984) was administered in Study 2 because it has been shown to be related to SES (Steen et al., 2020). The PSS assesses the degree to which these young adults perceive their lives as unpredictable, uncontrollable, and overloaded. Indeed, in our sample, lower subjective social status during childhood was associated with higher perceived stress in young adulthood (*r* = -0.36, *p* = 0.012).

### Experiment

In Study 1, we instructed participants to watch muted movies of their own choice, while the stream of task-irrelevant auditory tones played. In Study 2, we asked participants to watch a series of images and press the space bar as soon as each image appeared. In both studies, the presentation of the auditory stimuli was the same and described in detail below. Participants instructed to focus on the information on the screen and ignore the background sound. Prior to the experiment in Study 2, participants performed a flanker task that indexed inhibitory control (Eriksen & Eriksen, 1979).

#### Auditory stimuli

Participants were immersed in auditory oddball streams at 55 *dBA* via speakers containing standard or deviant tones in either a pitch ascending or descending condition. The stimulus in the ascending stream was the rapidly ascending pitch in frequency from 600 to 1400 Hz, i.e., 600 Hz, 800 Hz, 1000 Hz, 1200 Hz, and 1400 Hz. The stimulus in the descending stream was the rapidly descending pitch in frequency from 1400 to 600 Hz, i.e., 1400 Hz, 1200 Hz, 1000 Hz, 800 Hz, and 600 Hz. Each tone lasted 50 ms, and the inter-trial interval was 600 ms. The deviant in each stream was the last pitch: 1600 Hz instead of 1400 Hz in an ascending stream and 400 Hz instead of 600 Hz in a descending stream. Each block consisted of 144 standards and 36 deviants (80/20 ratio), lasting roughly 2.5 minutes. Both ascending and descending streams were repeated 9 times. In total, there were 324 deviants in each sound condition and 648 deviants in total for each participant. The first 10 trials in each block were standards (to establish the expectation), followed by a pseudorandom distribution of standards and deviants, avoiding consecutive deviants.

#### Visual stimuli

In Study 1, the visual stimuli were muted movies based on participants’ choices, following the standard auditory MMN paradigm. In Study 2, visual stimuli were images selected from the IAPS database (Lang et al., 1997). All selected images were within 2.2 standard deviations of the standard ratings for both valence and arousal levels. In addition, the valence ratings ranged from 1 to 3.8 for the selected negative images and from 4.5 to 5.5 for the selected neutral images. Each image was presented for a duration of 5000 ms (± 100 ms jittered) with an inter-stimulus interval of 1000 ms (± 100 ms jittered). Twenty images formed a block with different conditions: 1) all neutral image condition, 2) all negative image condition, and 3) dynamic image condition that randomly mixed 10 negative and 10 neutral images. The dynamic image condition was included because it has been shown to taxed more executive functioning (Hao et al., 2019). Each image sequence condition was repeated 3 times, and no image was repeated. Therefore, this is a design with a repeated within-subject factor of 3 image sequence condition (neutral vs. negative vs. dynamic) by 2 sound condition (ascending vs. descending) with 3 repeated blocks. In total, 18 blocks were presented in a pseudorandom order with the constraint that no same combination of image and sound conditions appeared sequentially. Figure 3 depicts the study procedure.

**Figure 3.**
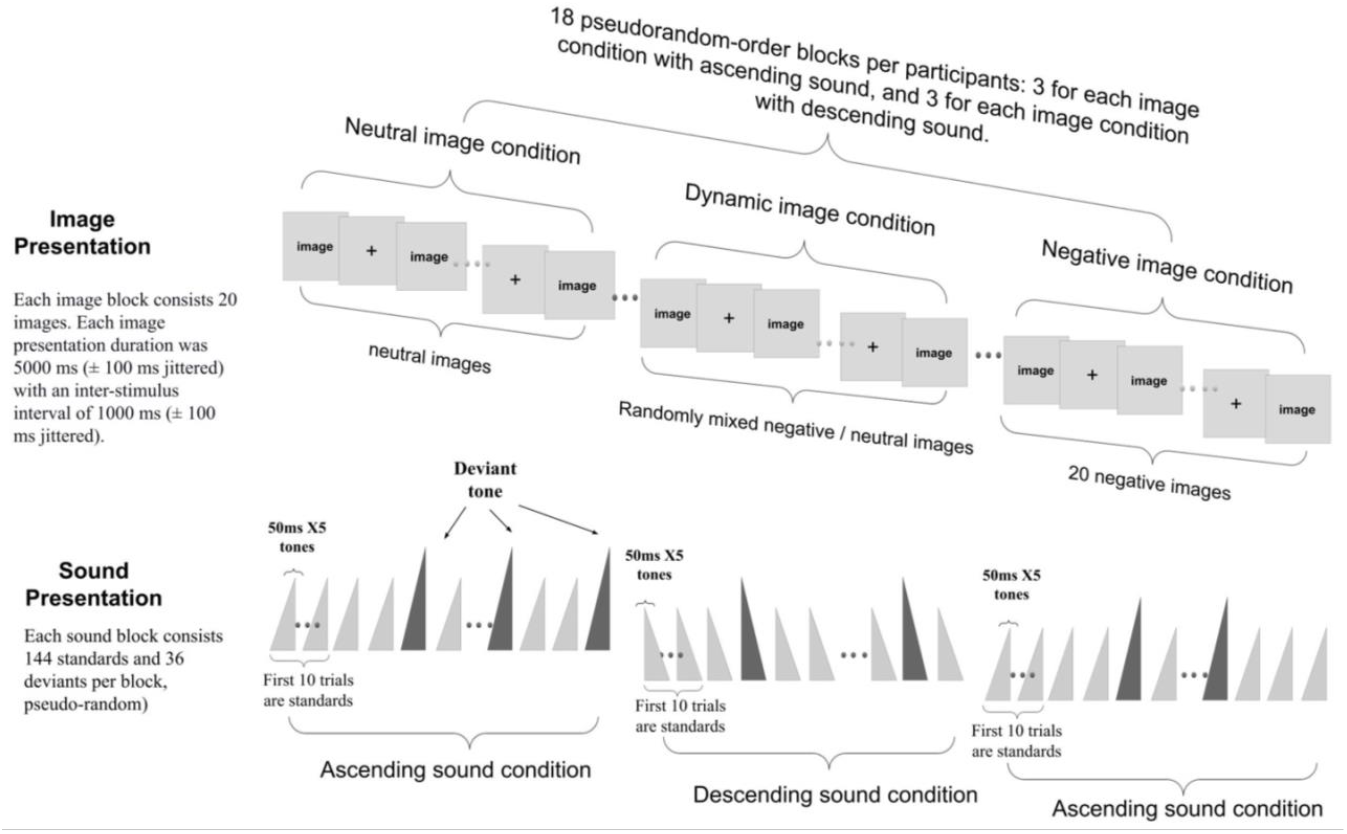
Study diagram

### EEG recording and processing

EEG data was recorded using a 128-channel BioSemi system at a sampling rate of 512 Hz. Audio and visual presentations, as well as key presses, were collected and synchronized with EEG. All EEG channels were offline referenced to the algebraic average of left and right mastoids and notch filtered (55∼65 Hz) to remove power-line noise. An ICA algorithm from EEGlab (Delorme & Makeig, 2004) was employed to detect and remove segments contaminated by eye movements, muscle, and cardiac artifacts. After removing the artifact components, the ICA source signals were transferred back to the original signal space for subsequent analysis.

For MMN analysis, the preprocessed EEG signal was further filtered within [0.5 20] Hz. Data were epoched into –100 ms to 350 ms trials, where time 0 was time-locked to the last tone of ascending or descending stimulus, which was 200 ms after the start of the complex tone pattern. The -300 ms to -200 ms period before the first tone of the trial was used for baseline correction. If the signal surpassed ±75 μV, the trial was excluded from further analysis. The MMN waveform was calculated as the difference between the ERP evoked by the deviant stimuli and the standard stimuli. Based on previous MMN literature, MMN waveforms from the frontocentral electrodes in the 100 ms to 250 ms time window were selected and averaged (Biosemi labeling: A1, C23, C21, C2, C11, D2, C24, see Figure 2A) to form a single MMN for subsequent analysis. We used the mean amplitude measures in this time window (Luck, 2014). We repeated the analyses with a single electrode located at the Fz site (Biosemi labeling: C21) to ensure the results were consistent. In addition, we analyzed P3a component (230 ms to 300 ms time window) at the frontal and parietal regions.

Emotional pictures that capture visual attention can increase the late positive potential (LPP) (Schupp et al., 2006). EEG responses were time-locked to the onset of each picture at 0 ms and extracted during the -300 ms to 4500 ms time window. We calculated grand average waveforms elicited by pictures at centroparietal pooling (Biosemi electrode labeling: A3, A4, A19, D16, B2, A5, A32) for each condition within each subject in the LPP ranges: the 400-600 ms and the 600 to 1,000 ms time window.

### Statistical Analyses

#### Analysis of ERP

To analyze the relationship between SES and MN, we used mixed effect models with each experimental condition nested in each participant as a random effect. Gender, age, and race were covaried in all models and all coefficients reported were standardized for comparability across measures. Mixed effect modeling was carried out with the package “lme4” (https://github.com/lme4/lme4/) in R. First, we reported the SES main effect on MMN separately for Study 1 and Study 2. Second, we explored the different relationships between SES and MMN within different contexts, such as sound frequency features, emotion and task types. We incorporated a 2-way interaction term of sound condition and SES in Study 1 and a 3-way interaction term of image condition, sound condition and SES in Study 2. To investigate whether the MMN relation to SES differ by task types, we combined two studies and analyzed the MMN relation to SES, incorporating an interaction term of Study type and SES. The same models were applied to P3a and LPP.

#### Analysis of reaction time to images

This analysis is specific to Study 2. We removed trails with reaction time (RT) greater than 1500ms and averaged the RT across the 3 blocks within each condition for each participant, leading to a total of 6 RT values per person. Log transformation was applied to RT due to its right skewness. The mixed linear model tested the main effects of SES, sound feature, image condition, and the 2-way and 3-way interactions among them, controlling for demographic covariates. Random effects were modeled for the sound feature and image condition nested within each participant. Additionally, we fitted another mixed linear model for log RT on MMN to examine whether the possible SES-related MMN differences were related to behavioral task performance. The same random effects and covariates were used as in the previous model. The majority of participants had missing values of at most 10% for each image condition that contains 60 trials. Removing 9 people with missing values over 10% for one or more conditions did not change our conclusions about SES relations to RT, so we reported the results including the whole Study 2 sample.

#### Analysis of flanker task

We examined the incongruent trials as they reflect executive attention and inhibitory control ability (Zelazo et al., 2014), focusing on response accuracy and reaction time separately. For accuracy, we employed linear regression to investigate the association between the accuracy rate among incongruent trials and SES, while controlling for gender and age. For reaction time (RT), we removed outlier trials (RTs below 100 ms or above 3 SD from each participant’s mean RT) using the criteria outlined in Zelazo et al. (2014), then computed and log-transformed the median RT for correct trials separately for congruent and incongruent trials. Linear regression was used, with log median RT as the response variable, SES as the primary predictor, and gender, age, race, and log median RT in congruent trials as covariates. To confirm the findings, we retested the model using the median RT differences between congruent and incongruent trials as the response variable. We also examined the relationship between MMN amplitude and flanker performance using the same random effects and covariates.

### Results

### Brain responses to changes in irrelevant sound are predicted by SES

At approximately 150 ms following the onset of the deviant sound, a distinct auditory evoked component of ERP is observed. The amplitude of the MMN is significantly correlated with SES in both Study 1 (*p* = 0.0004) and Study 2 (*p* = 0.0006), while controlling for demographic covariates and experimental conditions. In both studies, no significant main effects were observed for any of the covariates, while only the sound condition with ascending sound stream elicited greater amplitude (both studies *p* < 0.01). The complete statistical results are presented in Table 2.

**Table 2.** Statistics of MMN amplitude and its relation to SES as measured by subjective social standing and other predictors. Standardized beta coefficients (standard error) are reported for ease of interpreting effect sizes.

| | Study 1 ($N = 16$) | Study 2 ($N = 42$) |
| --- | --- | --- |
| MMN amplitude ($\mu V$) | -2.79 (0.29)<br>95% $CI = -3.35$ to $-2.22$ | -4.08 (0.25)<br>95% $CI = -4.61$ to $-3.63$ |
| SES | 0.58 (0.16) ***<br>95% $CI = 0.27$ to $0.89$ | 0.39 (0.11) ***<br>95% $CI = 0.17$ to $0.61$ |
| Sound condition<br>(*Low sound as reference*) | High: -0.89 (0.28) ** | High: -0.40 (0.08) *** |
| Image condition<br>(*Neutral image as reference*) | NA | Negative: -0.01 (0.10)<br>Dynamic: 0.07 (0.08) |
| Gender<br>(*Female as reference*) | Male: -0.01 (0.46) | Male: -0.16 (0.28) |
| Age | 0.06 (0.18) | -0.08 (0.13) |
| Race<br>(*Asian as reference*) | White: 0.27 (0.32)<br>African American: 1.22 (0.52) * | White: 0.06 (0.30)<br>African American: NA |
| Notation: *** $p < 0.001$, ** $p < 0.01$, * $p < 0.05$ | | |

We also conducted analyses with MMN from a single electrode located at Fz, and found that the results are consistent with the grand averaged MMN across electrodes around frontocentral region. Figure 4 illustrates the MMN waveforms for each participant and the relationship between MMN amplitude and SES, specifically for Study 2.

**Figure 4.**
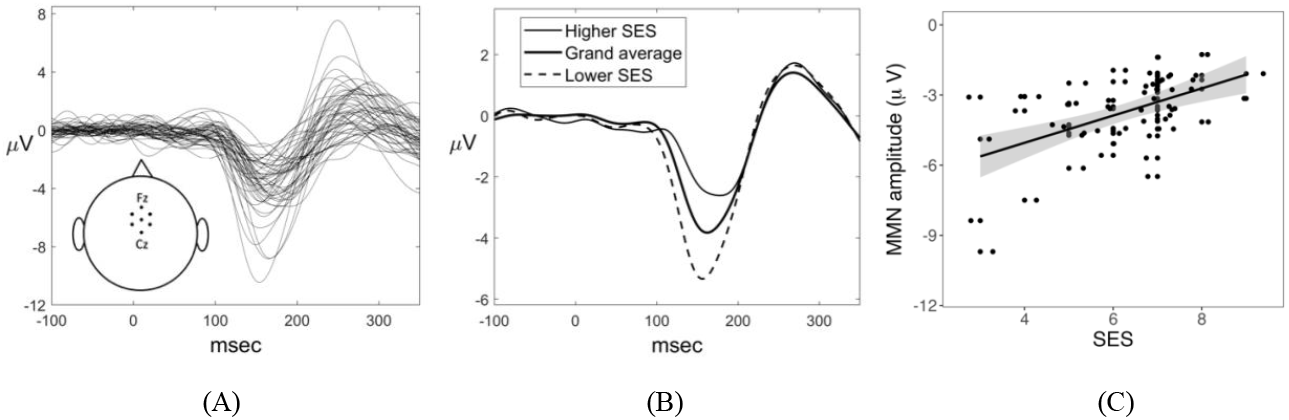
Visualization of MMN relation to SES. (A) Grand average of the MMN waveforms at the frontocentral region for each participant, averaged across experimental conditions. The waveforms are derived from the 7 electrodes surrounding Cz and Fz channels. (B) MMN waveforms for participants grouped according to their SES levels. To group participants, we identified individuals with SES scores greater than one standard deviation above the mean (SES > 7.5; 5 people) as the high-SES group and those with scores below one standard deviation from the mean (SES < 4.5; 8 people) as the low-SES group. The grouping is only for visualization purpose. (C) Scatter plot of MMN amplitude and SES for each individual.

To investigate whether the effect of SES on MMN varies across contexts, we tested the interaction of the experimental conditions with SES on MMN. However, none of the interactions were found to be significant (all *p* > 0.112). Additionally, controlling for perceived stress did not influence the SES effect.

Furthermore, we examined whether the MMN relation to SES differs across studies. Since the effects of SES were not affected by experimental conditions in either Study 1 or Study 2, we averaged MMN across conditions for each participant. The interaction of study type and SES is not significant (*p* = 0.6119), and the SES main effect hold (*β* = 0.53, *S*.*E*. = 0.11, 95% *CI* = 0.31 to 0.75, *p* < 0.0001), indicating that the MMN and SES relation does not vary by study type.

Additionally, despite that we observed clear P3a component at the frontal and parietal regions, indicating automatic orienting of attention to novelty in the context of unattended stimuli (Friedman et al., 2001), there is no SES differences on P3a in both regions (all p > 0.6) although we can see clear SES differences in MMN. Figure 5 illustrates the ERP waveforms and the relationship to SES in Study 2.

**Figure 5.**
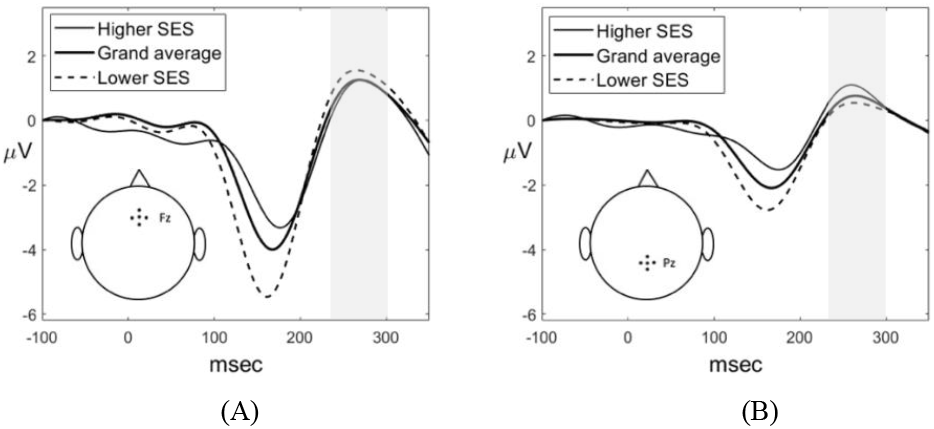
(A) P3a at the frontal regions. The waveforms are derived from the 5 electrodes surrounding Fz channels, including C12, C20, C21, C22 and C25. (B) P3a at the parietal region. The waveforms are derived from the 5 electrodes surrounding Pz channels, including A4, A5, A19, A20 and A32. The ERP waveforms are averaged across experimental conditions for each participant. To group participants, we identified individuals with SES scores greater than one standard deviation above the mean (SES > 7.5; 5 people) as the high-SES group and those with scores below one standard deviation from the mean (SES < 4.5; 8 people) as the low-SES group. The grouping is only for visualization purpose.

Overall, our findings suggest that lower SES is associated with greater brain detection to sound changes, and this effect is consistent across sound features, emotional contexts, and study types. Furthermore, lower SES individuals did not involuntary orient their attention to deviance more than their higher SES counterparts.

### Brain responses and behavioral performances in attended tasks are invariant across SES

To investigate the effect of SES on sustained attention, we analyzed LPP induced by emotional images in Study 2. We focused on the time ranges of 400-600ms and 600-1000ms, which are known to reflect sustained allocation of attention (Schupp et al., 2006). Our results show that unpleasant images elicited a significant increase in LPP (p < 0.0001), consistent with prior research. However, there is no significant relationship between LPP and SES in either time range (p = 0.778 and p = 0.691, respectively), nor is there an interaction between SES and image or sound conditions on LPP (all p > 0.14).

Furthermore, we found no significant relationship between SES and RT in any experimental condition (all p > 0.513). Although participants performed well in the image detection task across different conditions (mean RT = 423.74 ms), there was considerable variation in RT (range: 222.07 to 943.81 ms), indicating that the task was not too easy. Participants responded more slowly to negative images than neutral ones in both ascending (p = 0.029) and descending sound conditions (p = 0.008), indicating successful manipulation of emotional context (Pereira et al., 2010). Moreover, there was no significant interaction between condition and RT on MMN (all p > 0.171).

There was no significant difference in participants’ performance on the flanker test based on their SES. After controlling for demographic covariates, we found no association between accuracy in the incongruent condition and either SES or MMN (all *p* > 0.200). Additionally, after excluding outlier trials and controlling for demographic covariates and log RT on the congruent trials, we found no significant association between log RT on accurate incongruent trials and either SES or MMN (all *p* > 0.560). The use of differences in median RTs between congruent and incongruent as the response variable did not change any of these results (all *p* > 0.356).

In conclusion, our results suggest that lower-SES participants were as focused on the task as their higher-SES counterparts, as evidenced by the lack of significant differences in LPP and RT across SES. And in our sample, the inhibitory control as measured by flanker task is also invariant across SES.

## Discussion

We investigated how perceived childhood social standing relates to adult brain responses to irrelevant sound changes across two studies. Our novel findings suggest that lower perceived childhood social standing is associated with enhanced automatic change detection. Furthermore, our findings remained consistent across two studies that involved different attended tasks and various levels of salience of the unattended auditory deviance.

Previous research on SES effects on irrelevant information processing has mainly focused on effortful attentional control (D’Angiulli et al., 2008; D’Angiulli, Weinberg, et al., 2012; Li et al., 2022; Stevens et al., 2009, 2015), but our study provides a unique contribution by focusing on preattentive processing and demonstrating SES-related differences in this early stage of processing. We suggest that SES may also affect irrelevant information processing without altering attention allocation. The lack of SES differences in P3a magnitude suggests that lower-SES participants did not pay more attention, even involuntarily, to the unexpected change in ambient sound than their higher SES peers. The lower-SES participants’ brains might perceive the deviance as more salient than those of higher SES, but this did not reach a level to further impact the degree of attentional processing of the information. Our findings cannot be explained by extra attention to deviance in lower-SES participants.

Our study also examined unattended auditory information processing while participants focused on visual information, showing that the increased sensitivity to detecting changes in task-irrelevant auditory information did not interfere with participants’ visual attention processing, as supported by our LPP results. In general, affective stimuli can influence attention and compete with top-down control on perception (Vuilleumier, 2005), and processing emotional contexts with varying valence and arousal levels can tax executive functioning. However, in our study, we found that behavioral responses and brain responses to the visual task were similar across all SES groups, suggesting no difference in visual attention allocation. In addition, SES effect on MMN response to unattended auditory changes was consistent across study types (passively viewing in Study 1 and detection task in Study 2). However, we should be cautious when interpreting the absence of an interaction effect, as detecting significant interactions requires a larger sample size (Gelman et al., 2020).

Moreover, we used a subjective measure of SES instead of objective measures like maternal education or family income, which may provide a more nuanced gradient in measuring SES and capture the relationship between SES, health, and brain functions (Adler et al., 2000; Anwyl-Irvine et al., 2021; Hoebel et al., 2017; Singh-Manoux et al., 2005). Despite our participants being raised in relatively affluent families, their subjective social standing during childhood was related to their perceived stress in adulthood (c.f., Steen et al., 2020). We controlled for potential confounding variables, such as age, gender, race, allowing for a more rigorous examination of the link between SES and brain functions. By recruiting from a top US university, we reduced the variance in participants’ current objective SES and focused on the correlates of subjective childhood SES. Our findings revealed significant differences in processing irrelevant auditory information based on perceived childhood social standing.

One potential explanation for the link between lower subjective social status and stronger automatic change detection is that individuals who perceive themselves as lower in social status may be more attuned to deviances that signal threats and dangers, particularly in their social interactions. Prior research has found that individuals with lower subjective social status exhibit greater fear reactivity to lab-induced social stress, report social stress tasks to be more threatening, and display higher stress-induced inflammation than those with higher subjective social status, even after controlling for objective SES (Derry et al., 2013; Rahal et al., 2020). Neuroimaging studies have also revealed that lower subjective social status is associated with reduced gray matter volume in the perigenual areas of the anterior cingulate cortex, a brain region implicated in emotion regulation and stress response (Gianaros et al., 2007), as well as reduced total volume of the amygdala, another brain region involved in emotional processing that is susceptible to environmental stress (Assari et al., 2020).

In our study, the observed automatic change detection may be more salient for participants with lower subjective social status because they are accustomed to taking subtle changes in their environment more seriously. However, as participants were focused on the visual task and our task was not inherently threatening, the detected deviance was processed with higher priority only at the early auditory processing stage without entering consciousness to draw attention. As a result, a salience-related enhancement in MMN amplitude may occur without lower SES participants paying more attention to task-irrelevant sounds (c.f. Fitzgerald and Todd 2020; Quiroga-Martinez et al. 2020; Quiroga-Martinez et al. 2021).

In summary, our study suggests that lower-SES individuals may not necessarily be less able to inhibit distractions, as evidenced by their similar performance on the flanker task compared to higher-SES individuals. Instead, they may be more responsive to their environment and more likely to notice and react to subtle changes. While this may seem advantageous, it can also be costly as it reflects a more reactive coping style (Aron et al., 2012, pp.264). However, our study did not explore the potential behavioral consequences associated with this difference in cognitive processing between SES groups.

One limitation of our study is that we did not examine the relationship between current social status and childhood social status on MMN. However, we controlled for current perceived stress, which reflects an individual’s perception of stress in their life and can be influenced by factors such as financial strain, discrimination, and environmental stressors. Perceived stress can reflect both the experiences of young adults and their perceived control over these experiences, which are associated with childhood perceived social standing in our study. Another limitation is that our findings may not be generalizable to populations other than college students, and the full implications of these neural differences are yet to be understood.

The present study provides an essential first step by establishing a link between SES and the neural responses underlying change detection of irrelevant information processing. Our findings highlight that despite small differences in behavioral and neural responses to attended stimuli, there were significant neural effects related to perceived social status.

## Data availability statement

The dataset and code for the current study are available from the corresponding author on reasonable request.

